# Individualized Phenotyping of Functional ALS Pathology in Sensorimotor Cortex

**DOI:** 10.1101/2025.01.13.632667

**Authors:** Avinash Kalyani, Alicia Northall, Stefanie Schreiber, Jascha Brüggemann, Stefan Vielhaber, Marwa Al Dubai, Abrar Benramadan, Hendrik Mattern, Oliver Speck, Christoph Reichert, Esther Kuehn

**Affiliations:** Institute for Cognitive Neurology and Dementia Research (IKND), Otto-von-Guericke University Magdeburg, 39120, Germany; German Center for Neurodegenerative Diseases (DZNE), Magdeburg, 39120, Germany; Leibniz Institute for Neurobiology (LIN), Otto-von-Guericke University Magdeburg, Germany; Center for Behavioral Brain Sciences (CBBS) Magdeburg, Magdeburg, 39120, Germany; Clinic for Neurology, Otto-von-Guericke University Magdeburg, 39120, Germany; Department Biomedical Magnetic Resonance (BMMR), Otto-von-Guericke University Magdeburg, Germany; Research Campus STIMULATE, Otto von Guericke University, Magdeburg, Germany; Hertie Institute for Clinical Brain Research, 72076 Tübingen, Germany; German Center for Neurodegenerative Diseases (DZNE), Tübingen, Germany; Nuffield Department of Clinical Neurosciences, University of Oxford, United Kingdom

**Keywords:** Amyotrophic Lateral Sclerosis (ALS), PLSR, sensorimotor cortex, 7T-fMRI, disease progression

## Abstract

Amyotrophic Lateral Sclerosis (ALS) is a progressive neurodegenerative disease characterized by the loss of motor neurons in primary motor cortex (MI), leading to muscle weakness, atrophy, and death within a median of three years. Even though ALS is characterized by different disease subtypes affecting different body parts, individiualized phenotyping of functional ALS pathology has so far not been achieved. We recorded 7 Tesla functional MRI (7T-fMRI) data while ALS patients and matched controls moved affected and non-affected body parts in the MR scanner. We applied robust Shared Response Modeling (rSRM) for capturing ALS-specific shared responses for group classification, and Partial Least Squares (PLS) regression for relating the latent variables to clinical subtypes and the degree of disease progression. We show that both functional connectivity and functional activation in MI are a predictor for disease onset site. However, disease severity could best be predicted by functional connectivity rather than pure activation changes. Critically, we show that functionally disease-defining information in MI is not strongest in the area that is behaviorally first-affected, deviating from the behavioral phenotype of the patients. When computing the model’s weight distribution of the King stage classification and projecting them back into voxel space, the highest mean weights are present in the foot and tongue/face regions that seem to drive disease progress. Our data highlight the importance of 7T-fMRI task-based functional connectivity measures for classifying ALS-patients, and provide evidence that a single 7T-MRI scan can be used for identifying a disease signature of each individual ALS patient.

## 1. Introduction

Amyotrophic Lateral Sclerosis (ALS) is the most common form of motor neuron disease. It is characterized by progressive muscle weakness and atrophy, ultimately leading to death with a median survival time of three to five years from disease onset (Al-Chalabi et al., 2016; Norris et al., 1993). Both upper and lower motor neuron degeneration contribute to the classical ALS phenotype, while some patients present with a specific motor neuron phenotype. Behaviorally, ALS presents with a topographic disease phenotype, in which symptoms initially present focally in one body part. This onset is highly varied between patients, with roughly equal numbers experiencing onset in the head/face area (bulbar onset), the arms and hands (upper limb onset), or the legs and feet (lower limb onset) (Ravits and La Spada, 2009). According to the current dominant view, the disorder then progressively spreads to other body parts over the course of the disease, typically ending with bulbar and respiratory involvement (Chio et al., 2009). A functional signature of ALS in affected sensorimotor cortex, however, is so far lacking.

This topographic disease profile is partly reflected by microstructural alterations in the primary motor cortex (MI) that can be described in-vivo. Using 7T-MRI, a recent study showed that iron accumulation in the deeper layers of MI is highest in the area that represents the first affected body part (Northall et al., 2023a). However, other microstructural changes, such as calcium accumulation in upper layers, appear atopographic in MI (Northall et al., 2024). In addition, low-myelin borders that usually separate major body part representations in MI and hence are always located between two body part regions (Kuehn et al., 2017; Northall et al., 2023b), show early calcium accumulation and also degenerate early in ALS (Northall et al., 2024), which may speak against a strictly topographic disease profile. Despite its clinical significance, the individual functional topographic and/or atopograhic signature of ALS-patients has not been described in great detail. This is a critical limitation, as the functional signature of ALS in the sensorimotor cortex is expected to show a clear relationship to behavioral impairments and disease spread, and may more dynamically respond to treatments compared to structural measures. For instance, studies have shown that disruptions in functional connectivity can be detected before significant structural changes are observed in the brain (Filippi et al., 2020). This pattern has been noted in various neurodegenerative conditions, including Alzheimer’s disease and Parkinson’s disease, where early functional impairments can serve as indicators of subsequent structural degeneration (Dan et al., 2023; Ou et al., 2021). Understanding these functional signatures could allow for monitoring disease progression or the effectiveness of new medications with clear functional output metrics. However, to achieve this, we first need a clear understanding of the functional signature of ALS in the human sensorimotor system, which is so far lacking.

Recent advances in neuroimaging techniques, particularly 7T-fMRI, have provided new avenues for describing small-scale circuits of the sensorimotor system in living participants and patients (Kuehn and Pleger, 2020; Schreiber et al., 2021). Compared to standard 3T-fMRI, 7T-fMRI offers higher spatial and temporal resolution, providing images of functional brain networks and topographic activation that are more precise (Kreitz et al., 2023), which allows detecting subtle changes in brain activity that might otherwise be missed. Additionally, the shortened T2 relaxation rates at higher field strengths, such as 7T, improve the sensitivity of BOLD fMRI, making it especially beneficial for capturing fine-grained neural dynamics and subtle differences in activation patterns. These advantages make 7T-fMRI a powerful tool for studying neurodegenerative diseases like ALS, where early and localized changes in brain function are critical for understanding disease progression. Moreover, since ALS patients often have difficulties remaining still for extended time periods, the ability of 7T-fMRI to obtain high-quality images in a shorter amount of time is a significant advantage, making it a superior choice for studying the intricate neural correlates of ALS. Nevertheless, few studies have employed 7T-MRI for studying MI pathology in a group of ALS patients, and most are confined to structural measures (Cosottini et al., 2016; De Reuck et al., 2017; Johns et al., 2019; Northall et al., 2023a). The few functional 7T-MRI studies that exist only measured resting state activation but did not probe the sensorimotor cortex when in action (Barry et al., 2021), and leave it open which functional variables best reflect disease progression (Fox and Greicius, 2010).

7T-fMRI patient studies come with their own set of challenges. One key challenge is to extract ultra-high resolution functional time series at the single patient level that represents the mesoscale architecture of the task representations but to nevertheless conduct group statistics to compare a single patient’s profile to average functional patterns, as well as to clinical variables. Such computations require advanced statistical and computational methods that allow preserved resolution at the single patient level, in addition to comparing extracted functional patterns across patients. To address these challenges and to take full advantage of the ultra-high-resolution images, we aimed to employ techniques that can effectively handle high-dimensional fMRI datasets, namely robust Shared Response Modelling (rSRM) (Kalyani et al., 2023; Turek et al., 2018) and Partial Least Square Regression (PLSR) (Krishnan et al., 2011; McIntosh and Lobaugh, 2004). We used the former (i.e. rSRM) to capture group specific shared neural responses for classifying ALS vs control participants, and the latter (i.e. PLSR) to identify the functional architecture of ALS that is associated with disease progression.

rSRM is a multivariate method designed to capture shared neural patterns across subjects, even in the presence of individual variability (Turek et al., 2018). Unlike traditional alignment approaches, rSRM identifies a low-dimensional shared space that represents common neural responses, while discarding individual-specific components, such as noise or anatomical differences. This makes rSRM particularly suited for ultra-high field (7T) fMRI data, where inter-subject variability can obscure shared functional signatures. By aligning neural responses across groups, rSRM enables direct comparisons of functional architectures, such as those observed in ALS patients versus controls, and enhances the detection of group-specific patterns relevant to disease classification.

PLSR, on the other hand, is a multivariate regression method suited for high-dimensional data that has been widely used across disciplines such as social science, bioinformatics and neuroscience (Krishnan et al., 2011; Mehmood et al., 2012). PLSR projects the functional response and the covariables (i.e., measures of disease progression) into a new space formed by its latent variables. To target the question of the a/topographic profile of functional markers, we used functional activation triggered by body part movements (hands, feet, lip/tongue) as well as the associated network centrality (i.e, Eigenvector centrality mapping (ECM), (Lohmann et al., 2018)), while, we used different markers of ALS disease progression measured by neurologists (i.e., ALSFRS, PUMNS) as covariables. In addition, we tracked functional changes of a subset of patients over time. Given PLSR can handle high-dimensional data, it is an ideal method for extracting relevant features from complex 7T-fMRI data, and for computing relationships between feature dimensions and behavioral as well as clinical profiles (Krishnan et al., 2011; McIntosh and Lobaugh, 2004; Van Roon et al., 2014). Combining these approaches provides valuable insights into the functional signature of the disease across different topographic locations and disease stages.

Specifically, using 7T-fMRI, we tested if (1) participants can be successfully classified into ALS-patients versus controls based on their functional profile of the sensorimotor cortex recorded during body part movements, (2) there are latent variables in the functional time series that reflect disease severity and site of onset, and if so, if they become most evident in functional activation or functional connectivity changes, and (3) disease-defining functional information is specific to the first-affected location in the sensorimotor cortex (i.e., topographic, reflecting the iron accumulation) or not specific to the first-affected location in the sensorimotor cortex (i.e., atopographic, reflecting the calcium accumulation and the early affected low-myelin borders), and whether there is any location in MI that is highly predictive of the severity of the disorder irrespective of onset site. The results of our study contribute to a deeper understanding of the neural mechanisms underlying ALS and offer potential functional neuronal signatures that characterize the disease for future clinical applications.

## 2. Methods

An overview of the methodologies applied and the experimental approach is depicted in **Figure 1**.

**Figure 1.**
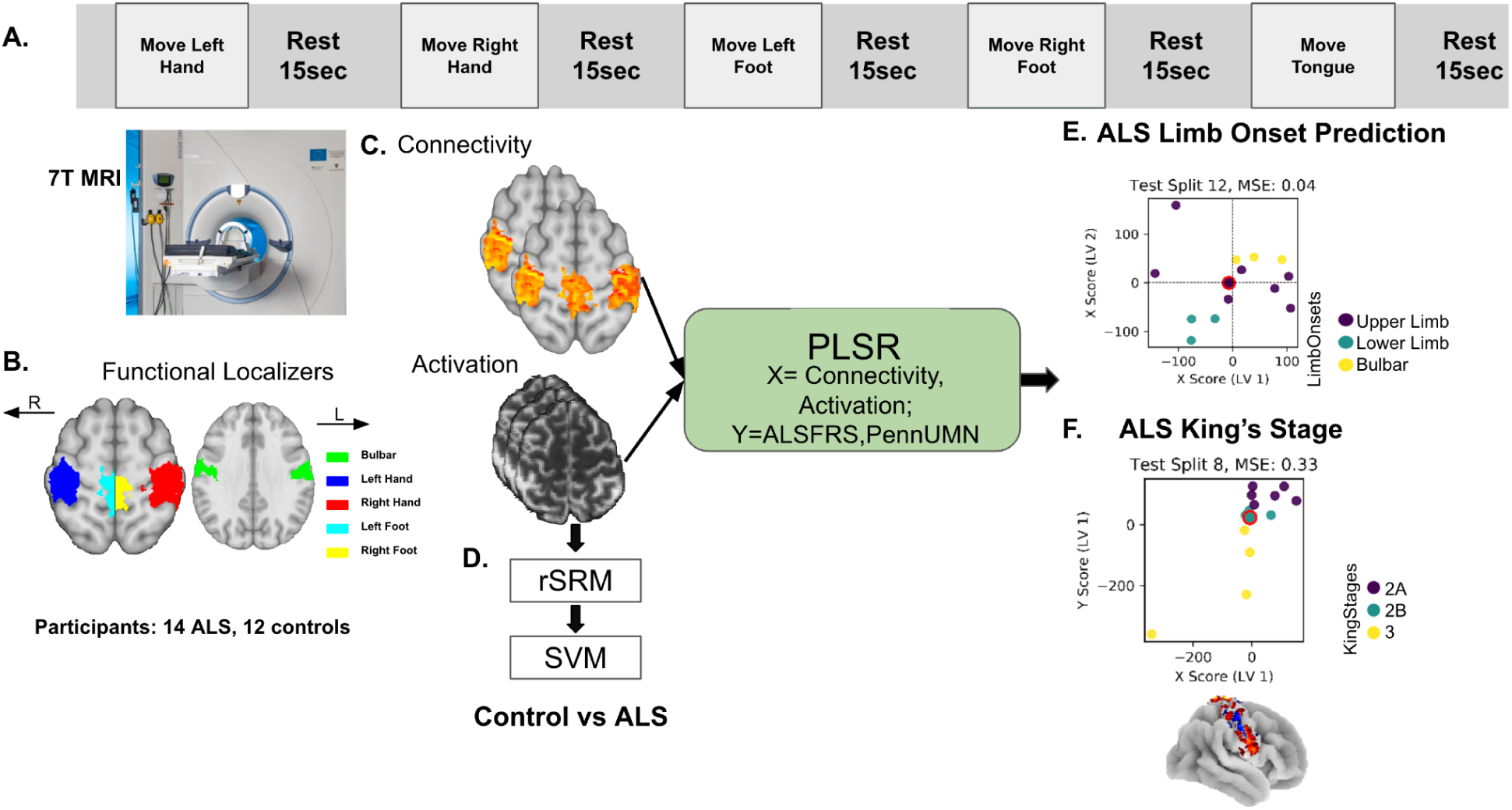
Experimental Design and Methodology. **(A)** During 7T-MRI scanning, participants and patients with a diagnosis of ALS were asked to move different body parts (left hand, right hand, left foot, right foot, and tongue/lip), each movement lasting 12 seconds, followed by a 15 seconds rest period. **(B)** Functional localizers identify sensorimotor cortices activated by these movements: red for right hand, blue for left hand, cyan for left foot, yellow for right foot, green for tongue/face. **(C)** Connectivity and activation maps are generated from 7T-fMRI data, highlighting brain regions involved in the tasks. **(D)** Classification of ALS patients versus controls using robust Shared Response Modeling (rSRM) and Support Vector Machines (SVM) classification obtained from functional patterns. **(E)** Prediction of ALS onset site is achieved using PLSR, with connectivity and activation as predictors (X) and ALSFRS and PUMNS as outcomes (Y). The scatter plot illustrates the latent variables (LV1 and LV2) for an example subject, showing clustering based on onset type (upper limb, lower limb, bulbar), with an MSE of 0.04. **(F)** ALS King’s stage prediction is similarly conducted using PLSR, with scatter plots depicting the relationship between the X and Y scores in the latent variable space, highlighting the clustering of subjects according to King’s stages (2A, 2B, and 3), with an MSE of 0.33.

### 2.1 Participants

We acquired fMRI data from n=14 ALS patients (6 females, age: mean = 56.07 years, SD = 15.62 years) and n=12 matched healthy controls (6 females, age: mean = 61.1 years, SD = 11.9 years). The patients were recruited from the University Hospital Magdeburg, Jena and Dresden, and participated in the study between June 2018 and January 2024. The patients were first scanned after their first diagnosis of ALS. Controls were individually matched to a subset of 12 of the patients based on age (±2 years; t(22) = −0.15, P = 0.883), handedness, gender, and years of education (±4 years; patients: mean = 14.5 years, SD = 2.7 years; controls: mean = 15.4 years, SD = 2.7 years; t(22) = −0.82, P = 0.422). For two patients, no matched controls could be measured due to a software update that occurred within the time of data collection. Participants and patients underwent one 7T-MRI scan as well as behavioral assessments. Clinical assessments were performed by an experienced neurologist with the patients only. Among the patients, n=8 had upper limb onset, n=3 had lower limb onset, and n=3 had a bulbar onset (considered bilateral, see **Table 1**). n=3 patients were scanned a second (and third) time to allow tracking of disease progression (P1: after 5 months, P6: after 7 months, P4: after 8 months, after 2 years and 4 months). Healthy matched controls were recruited from the DZNE database in Magdeburg, Germany, with exclusion criteria including sensorimotor deficits, neurological disorders, and contraindications for 7T MRI. All participants and patients provided written informed consent prior to being scanned and were compensated for their participation. The study was approved by the local Ethics Committee of the Medical Faculty of the University of Magdeburg.

**Table 1.**
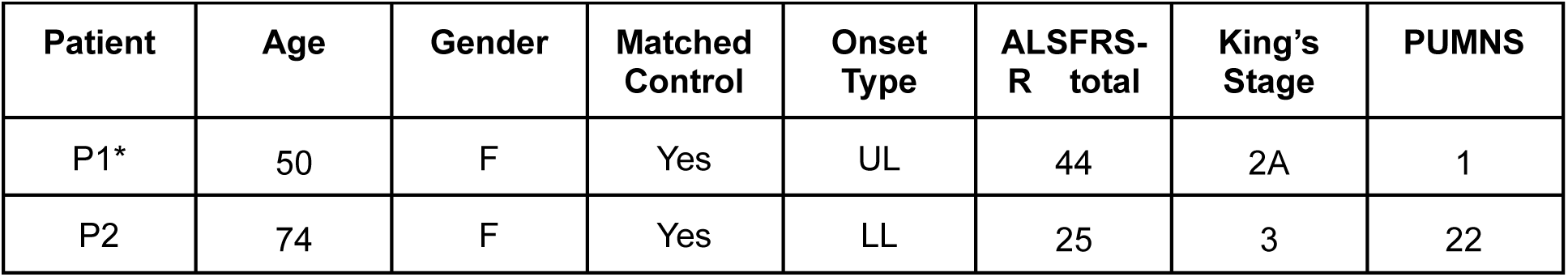
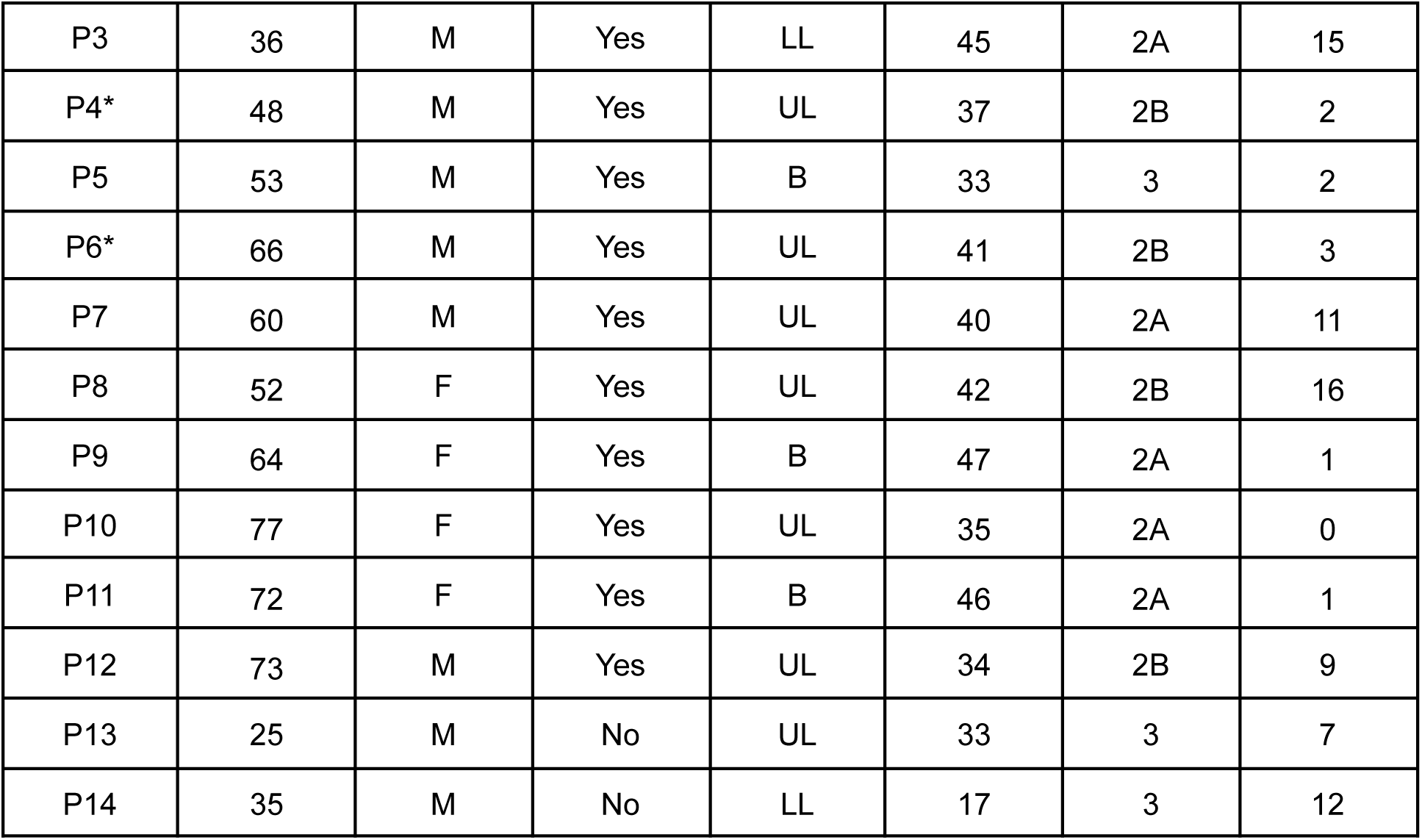
Clinical and demographic information on ALS patients. Onset type reflects upper limb (UL), lower limb (LL), or bulbar (B) onset site. The ALS Functional Rating Scale-Revised (ALSFRS-R) indicates disease severity, where lower values indicate greater impairment, with subscores for fine, gross, bulbar motor and respiratory function. The King’s stage indicates the stage of disease progression based on the ALSFRS-R score, where stage 2A reflects the involvement of one body part and that a clinical diagnosis has taken place, while stage 2B and stage 3 reflect the subsequent involvement of second and third body parts, respectively. The Penn Upper Motor Neuron Scale (PUMNS) score indicates clinical signs of upper motor neuron involvement, with higher scores indicating greater impairment (mean = 6.92, SD = 7.46, range: 0–22). F = female, M = male, P = Patient, * Patients scanned more than once

### 2.2 7T MRI Data Acquisition

MRI data was collected using a 7T MRI scanner (Siemens Healthineers) equipped with a 32-channel head coil (Nova Medical Inc.) at Otto-von-Guericke University, Magdeburg, Germany. In a single 7T-MRI session, we acquired whole-brain MP2RAGE images with a spatial resolution of 0.7 mm isotropic (sagittal slices), with a repetition time (TR) of 4800 ms, echo time (TE) of 2.01 ms, field of view (FOV) read of 224 mm, flip angles of 5°/3°, inversion times of 900/2750 ms, and a bandwidth of 250 Hz/Px. Additionally, whole-brain functional images were obtained with a spatial resolution of 1.5 mm isotropic using an EPI gradient-echo blood oxygen level-dependent (BOLD) sequence comprising 81 slices, with a TR of 2000 ms, TE of 25 ms, FOV read of 212 mm, interleaved acquisition, GRAPPA factor of 2, and a Simultaneous Multi-Slice (SMS) factor of 2.

Functional imaging encompassed a blocked design paradigm, comprising 12-second periods of body part movements (left/right foot: moving the toes, left/right hand: tapping the fingers, tongue/face: moving the tongue against the lip while having the mouth closed) interspersed with 15-second rest intervals (see Northall et al., 2023a for more details). Participants underwent pre-scan training to familiarize themselves with the movements, and visual cues were provided within the scanner (grey background, black color) to prompt specific movements (e.g., ‘prepare left hand’, ‘move left hand’). Each movement was repeated four times, resulting in a single run of a total 20 trials. Fingerless braces were worn to minimize hand movements during scanning. We also acquired whole-brain, 0.5 mm isotropic resolution susceptibility-weighted (SWI) images in 10/14 patients (transversal slices, repetition time = 22 ms, echo time = 9 ms, field of view read = 192 mm, flip angle = 10°, bandwidth = 160 Hz/Px), as two patients could not be scanned any longer due to fatigue and discomfort (note that the SWI sequence was measured last). The total duration of the scanning session was approximately 75 minutes (Northall et al., 2024). The structural data of this study are published in (Northall et al., 2024).

### 2.3 Clinical Assessments

The ALS Functional Rating Scale-Revised (ALSFRS) is a widely used tool to measure the severity of ALS, consisting of subscores for fine motor function, gross motor function, bulbar and respiratory function (Cedarbaum et al., 1999). Lower total scores on the ALSFRS indicate greater disease severity and functional impairment. On average, the ALSFRS score for the patients in this study was 41.1 (SD = 5.7, range: 25–47).

The King’s College (KC) Staging System categorizes the progression of ALS based on the ALSFRS score, offering a structured way to assess the involvement of different body regions over time. Stage 2A indicates the involvement of one body part, stage 2B reflects the involvement of a second body part, and stage 3 indicates the involvement of a third body part (Al-Chalabi et al., 2021). In this study, this categorization is referred to as *King’s Stage*.

The PUMNS is a clinical measure used to assess signs of upper motor neuron involvement in ALS patients. It quantifies the degree of upper motor neuron impairment, with higher scores indicating greater impairment (Quinn et al., 2020). In this study, the PUMNS ranged from 0 to 22, with a mean score of 6.92 and a standard deviation (SD) of 7.46. This wide range reflects the variability in upper motor neuron involvement among the participants.

### 2.4 Preprocessing of fMRI data

The preprocessing, conducted in Statistical Parametric Mapping 12 (SPM12), included motion correction, smoothing with a 2mm full-width at half maximum (FWHM) kernel and slice-time correction (Friston, 2003). Co-registration with anatomical 7T MP2RAGE image was executed in ITK-SNAP (v3.8.0) (Yushkevich et al., 2016), with manual adjustments based on anatomical landmarks as needed. For group-level analysis, we registered all subjects’ anatomical and co-registered functional data to MNI space after skull-stripping using ANTs’ standard ‘antsRegistrationSyN.sh’ script (Avants et al., n.d.). This script performs rigid, affine, and SyN registrations to the MNI 152 template (Fonov et al., 2011).

### 2.5 Statistical Analysis

2.5.1. **Functional activation**

Prior to statistical analysis, we subjected the fMRI time series to band-pass filtering within the frequency range of 0.01-0.1 Hz. In SPM12, a first-level General Linear Model (GLM) was then applied to produce t-statistic maps for each body part involved in the task (for example left hand, right hand, left foot, right foot, and tongue/face). This process creates subject-specific activations for cortical fields relevant to the disease, i.e., the left hand, the right hand, the left foot, the right foot, and the face area (Northall et al., 2023a, 2023b). These activation maps were then used as predictors in the model to categorize disease stages.

#### 2.5.2. Generating functional localizer masks within the sensorimotor cortex

Further group-level analysis involved generating localizer masks from the registered task fMRI data to isolate the most significant 1500 voxels for each distinct body part using the t-maps generated from the GLM analysis. This approach ensured that all ROIs had the same size, and patients who performed movements with reduced speed and/or strength compared to the controls were not disadvantaged in the analysis. For combining masks across subjects, a more stringent approach was applied: masks for each body part were intersected across all subjects, retaining only voxels that were consistently active in every subject. The resulting combined mask was then refined by identifying the largest connected cluster within this intersection using a connected component analysis. This method ensured that only the most robust and consistently activated regions were included for each body part (hand, tongue/face, and foot), focusing on areas that are most relevant to disease progression.

#### 2.5.3. Functional connectivity analyses

For functional connectivity analyses, we used Eigenvector Centrality Mapping (ECM), which is an analysis technique that measures the influence of a brain region within the entire network by considering both the number and quality of its connections. In the context of neuroimaging, ECM identifies key brain regions that act as hubs in neural communication pathways, and may likewise identify areas of reduced functional communication that characterize affected areas in MI in ALS patients. This method assigns higher centrality values to regions that are extensively connected to other well-connected regions, thus highlighting areas critical for information flow and integration across the brain (Lohmann et al., 2023). Utilizing Lipsia (Lohmann et al., 2001), we generated EC maps to visualize and quantify the centrality of each voxel within the brain network, defined by the combined functional localizer masks for all the movements (left and right hand, foot, and tongue/face), facilitating the identification of neural hubs that may be crucial for disease progression. The EC map was used as the second predictor for categorizing the disease stages (besides the activity profile) to test which of the two reflect better disease severity and site of onset (see hypothesis (2)).

In addition, seed-based functional connectivity was assessed across functionally localized regions by selecting seeds based on t-maps specific to task conditions. Each seed, defined by a radius of 5 voxels, provides averaged time series signals, and a dot product correlation was performed by correlating these signals with the time series from other voxels within the combined localizer mask. The resulting correlation values were then used to create a correlation map, which was employed in the subsequent analysis. To evaluate changes in functional connectivity across different stages of ALS, we computed the differences in the dot product correlation values between ALS patients and control subjects, averaging the results over all subjects to obtain a distribution of connectivity changes across various body parts and disease stages.

#### 2.5.4. Robust Shared Response Modeling (rSRM) for ALS vs. control classification

rSRM is a data-driven method designed to capture shared patterns of neural activity across multiple subjects, even in the presence of inter-subject variability. Unlike traditional approaches that may be affected by differences in brain anatomy or individual cognitive strategies, rSRM identifies a common low-dimensional shared space that aligns brain responses across participants. This method is particularly useful for fMRI studies, where variations in brain anatomy, noise, and subject-specific factors can obscure the detection of common neural signatures. By focusing on group-level shared information, rSRM improves the robustness and sensitivity of analyses, making it well-suited for tasks like group classification, especially in populations with high variability, such as patients with neurodegenerative diseases like ALS.

To distinguish between ALS patients and control subjects, we employed rSRM, creating two separate models to capture group-specific shared information. We refer to these models as rSRM_1 and rSRM_2, corresponding to the ALS and control groups, respectively. Our objective was to establish distinct shared spaces that encapsulate the unique neural activity patterns for each group. We trained rSRM_1 using functional data from the ALS group and rSRM_2 using data from the control group, ensuring that each shared space was optimized to capture the group-specific functional signatures in a leave-one-out cross validation manner. During training, we utilized task-based fMRI data masked for specific regions (e.g., hand, foot, tongue/face) to create the shared spaces.

For each test subject, the functional data was split into two parts after rearranging the fMRI time series to ensure synchronization across all subjects. The first half of the functional time series was treated as Run 1, and the second half as Run 2. Using the training data from these runs, we estimated the shared basis and individual components separately for both rSRM_1 and rSRM_2. The individual term was then discarded to focus solely on the shared group-specific features, thereby reducing individual variability and emphasizing shared patterns that are more likely to generalize across subjects (as shown in (Kalyani et al., 2023)).

Next, the test subject’s test data was projected into both the ALS and control shared spaces using the basis learned during training. This resulted in two sets of projected data—one corresponding to the ALS group and the other to the control group. These projected datasets were concatenated and served as input features for the classifier. By transforming the test data into both shared spaces, we ensured that the classifier could leverage group-specific information to make predictions about whether a subject belonged to the ALS or control group. In the classification step, leave-one-subject-out cross-validation was employed, with the subject’s group (ALS or control) serving as the class label. This process was repeated for all subjects in both groups, enabling the classifier to comprehensively learn patterns specific to each group while accounting for inter-subject variability. By discarding the individual term and focusing on the shared group-level information, our approach effectively utilized the distinct shared neural patterns in ALS and control subjects. This allowed us to improve classification performance by minimizing individual noise and emphasizing disease-specific features. For this binary classification task, the chance accuracy was set to 0.5, given the two distinct groups involved.

#### 2.5.5. Partial Least Squares Regression (PLSR) analysis

In order to test whether there are latent variables in the functional time series that reflect disease severity and/or disease onset, and if so, whether they become more evident in functional activation patterns and/or functional connectivity changes, we used PLSR and computed the following steps: PLSR projects the response and the covariables into a new space formed by its latent variables by decomposing both X matrix (i.e. task activation, connectivity) and y (i.e. ALSFRS, UMN-Penn scores) of *n (= 14)* number of subjects into orthogonal scores and loadings. The general formulation is given by :

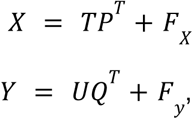

where, *X* = (*x*_1_ …. *x_n_*)*^T^*, represents the predictor matrix comprising of fMRI time series data of *n* subjects; *X* = (*y*_1_ …. *y_n_*)*^T^*, represents the response matrix i.e. behavioral data (in our case it is ALSFRS scores and UMN-Penn scores). *T* = (*t*_1_, …. *t_L_*) and *U* = (*u*_1_, …. *u_L_*) are the (n × L) matrices of L latent components corresponding to X and Y, respectively. The (p × L) matrix P and the (q × L) matrix Q are loadings and the (n × p) matrix Fx and the (n × q) matrix Fy are the matrices of residuals. Since our objective is to perform least squares regression in a low-dimensional latent space, the underlying assumption is that the latent component ui can be well predicted from ti from a relation such as:

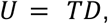

where, D is the (L × L) matrix. We need to maximize the covariance between *u_i_* and *u_i_* to satisfy the above assumption which produces weight matrices *W* = (*w*_1_, …. *w_L_*), and *c* = (*c*_1, …_*c_L_*) the equation is given by :

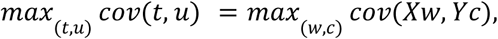

Our first aim was to relate functional activation and connectivity patterns to clinical outcomes. Our second aim was to group patients according to their disease progression (i.e., King’s Stages 2A, 2B, and 3), and the initial site of symptom onset (i.e., upper limb, lower limb, and bulbar regions). To achieve these goals, we employed two distinct PLSR models in a leave-one-out (LOO) cross-validation manner, each tailored to different predictive tasks: estimating ALSFRS scores and PUMNS from ECM and functional activation.

We utilized ECM values derived from preprocessed fMRI time series data as the independent variable (X), and the ALSFRS scores obtained from the patient cohort as the dependent variable (Y) (see **Figure 1F**). For the PUMNS predictions, we extracted fMRI time series data from the combined functionally localized masks specific to hand (i.e. both left and right hand regions), foot (i.e. both left and right foot regions), and tongue movements (i.e. bulbar region) together as the independent variable (X) and the PUMNS scores as dependent variable (Y) (see **Figure 1E**).

We evaluated the model’s performance by calculating MSE for each LOO cross-validation split. This assessment allowed us to project the test subject data onto the latent-variable space, facilitating an analysis of data clustering in relation to disease progression and symptom onset. For hypothesis (2), we used the response matrix Y, which included ALSFRS scores to predict the disease stage of ALS patients and UMN-Penn scores to predict the disease onset. For hypothesis (3), we projected the weights back into voxel space to determine if the weight distribution was topographic or atopographic. This was assessed by applying a paired t-test to the observed weights.

#### 2.5.6. Percent signal change over time

As an additional control analysis, we tested whether or not the strength of the recorded brain response reduces with disease progression. We calculated percent signal change for the subset of ALS patients (n=3) who were measured more than once. Percent signal change was calculated using the rest condition averaged over the whole brain within the two activity blocks as a baseline, and the task block as the parameter. This provides further information if brain responsivity can be used to predict disease progression (data shown in **Supplementary Material S1**).

To mitigate the inherent variability in BOLD signal interpretation, we z-scored the voxel-level time series and applied band-pass filtering (0.01 – 0.1 Hz) before calculating percent signal change. These preprocessing steps standardize the data, reduce noise, and focus on task-relevant neural activity. However, it is important to note that BOLD remains a relative measure influenced by physiological and vascular factors, such as cerebrovascular reactivity and cerebral blood flow, which may still impact the interpretation of longitudinal changes.

## 3. Results

### 3.1 Successful classification of participants into patients versus controls based on functional activation in sensorimotor cortex

To investigate if (1) participants can be successfully classified into patients and controls based on their functional activation in the sensorimotor cortex, we extracted shared response patterns across the group using rSRM, and trained SVM for predicting group membership (i.e., patient, control). Successful group classification could be reached, with an overall classification accuracy of 0.91±0.26. To test if the classification is mainly based on the region of the first-affected body part, or on regions of behaviorally not yet affected or later affected body parts, we compared classification accuracies between first-affected and not-first-affected body parts. Results revealed that first affected regions show significantly lower classification accuracy as compared to non-first affected regions (first-affected: 0.86 ± 0.13, non-first affected: 0.96 ± 0.23, (t(143) = -3.92, p = 1.50e-04, see **Figure 2B**).

**Figure 2.**
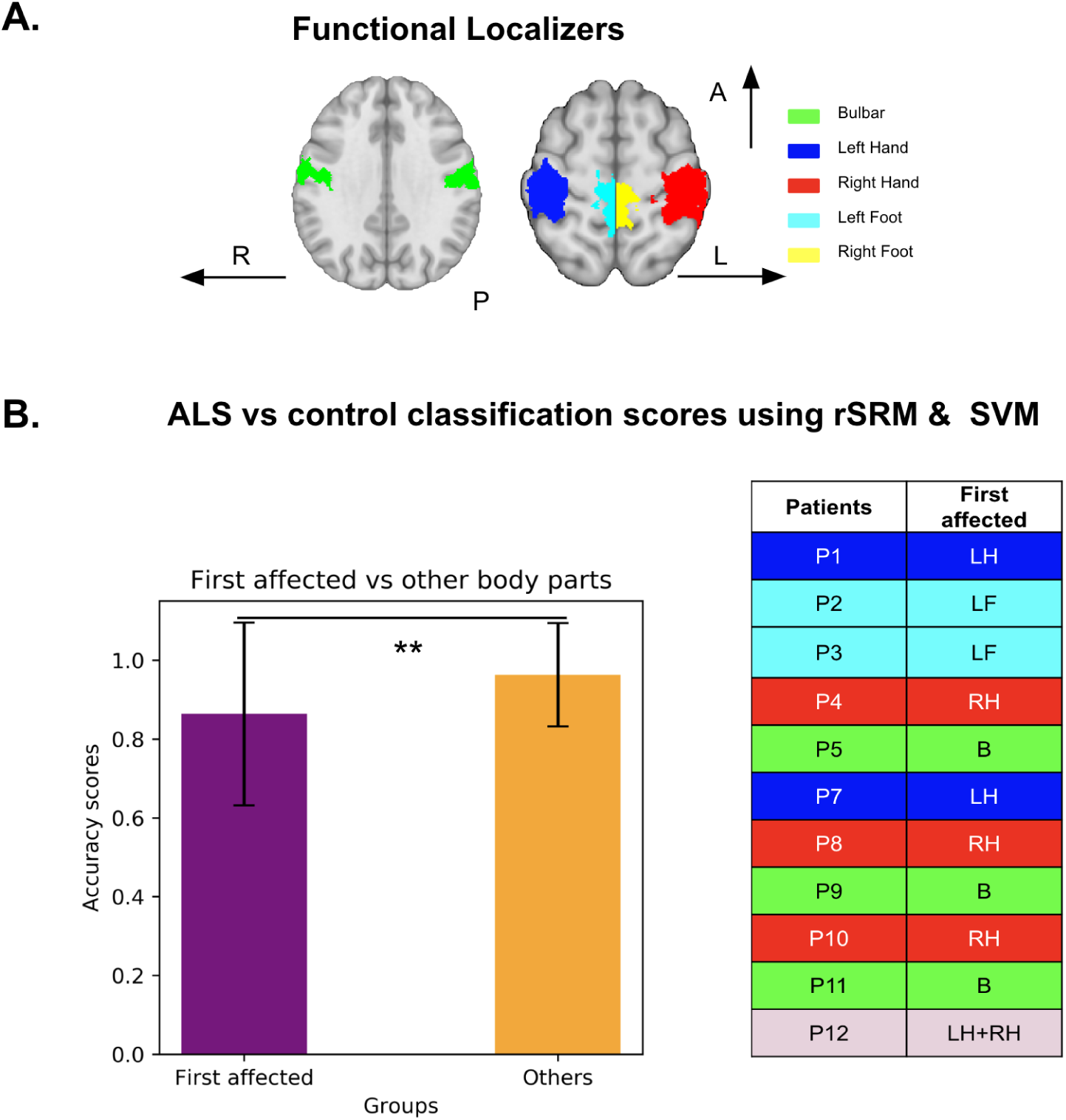
Classification of Participants into Patients versus Controls based on functional activation in sensorimotor cortex: **(A)** Axial brain slices display regions-of-interest (ROIs) used for functional localization. Different colors indicate body parts: blue for left hand (LH), red for right hand (RH), light blue for left foot (LF), yellow for right foot (RF), green for tongue/face (bulbar region - B). **(B)** Bar plot show classification accuracy scores for ALS patients versus controls using rSRM and SVM models. Scores are grouped by data extracted from the first affected body part (purple) versus non-first-affected body parts (orange); the first affected body parts show significantly lower classification accuracy (p < 0.01). **Table** lists the first affected body parts for each patient.

### 3.2 Identification of latent variables that reflect disease onset and severity

To test if (2) PLSR allows the identification of functional signatures that reflect disease onset and/or severity, and if so, if functional activation or functional connectivity changes are more sensitive to these clinical variables, we implemented PLSR using the functional activation and connectivity profiles as predictors, and related those to ALSFRS and PUMNS reflecting disease severity and onset. A combined localizer mask was used to derive data from different regions in MI (hand, foot, tongue), and the absolute values of averaged weights for these regions were plotted to assess their contribution to the prediction.

#### 3.2.1 Disease onset prediction

For functional activation, the overall average MSE was 1.02 ± 0.39 (mean ± SEM). The scatter plots in **Figure 3A** depict the separation between ALS onset types based on functional activation (purple: upper limb onset, cyan: lower limb onset, yellow: bulbar onset) across six test splits. While LV1 successfully separates lower limb onset from bulbar onset, upper limb onset cannot clearly be identified. This is reflected in the results when calculating the classification for each topographic area separately: the tongue/face region (bulbar) shows the highest average weight (5.3 ± 0.1e-5), followed by the foot region (3.8 ± 0.1e-5) and the hand region with lowest weights (1.9 ± 0.1e-5, see **Figure 3**). The histogram in **Figure 3C** shows the distribution of MSE values across splits, with the majority of splits clustered below 3, indicating good predictive performance. However, a wider spread of MSE values, with some splits showing MSEs as high as 6, suggests more variability in model performance.

**Figure 3:**
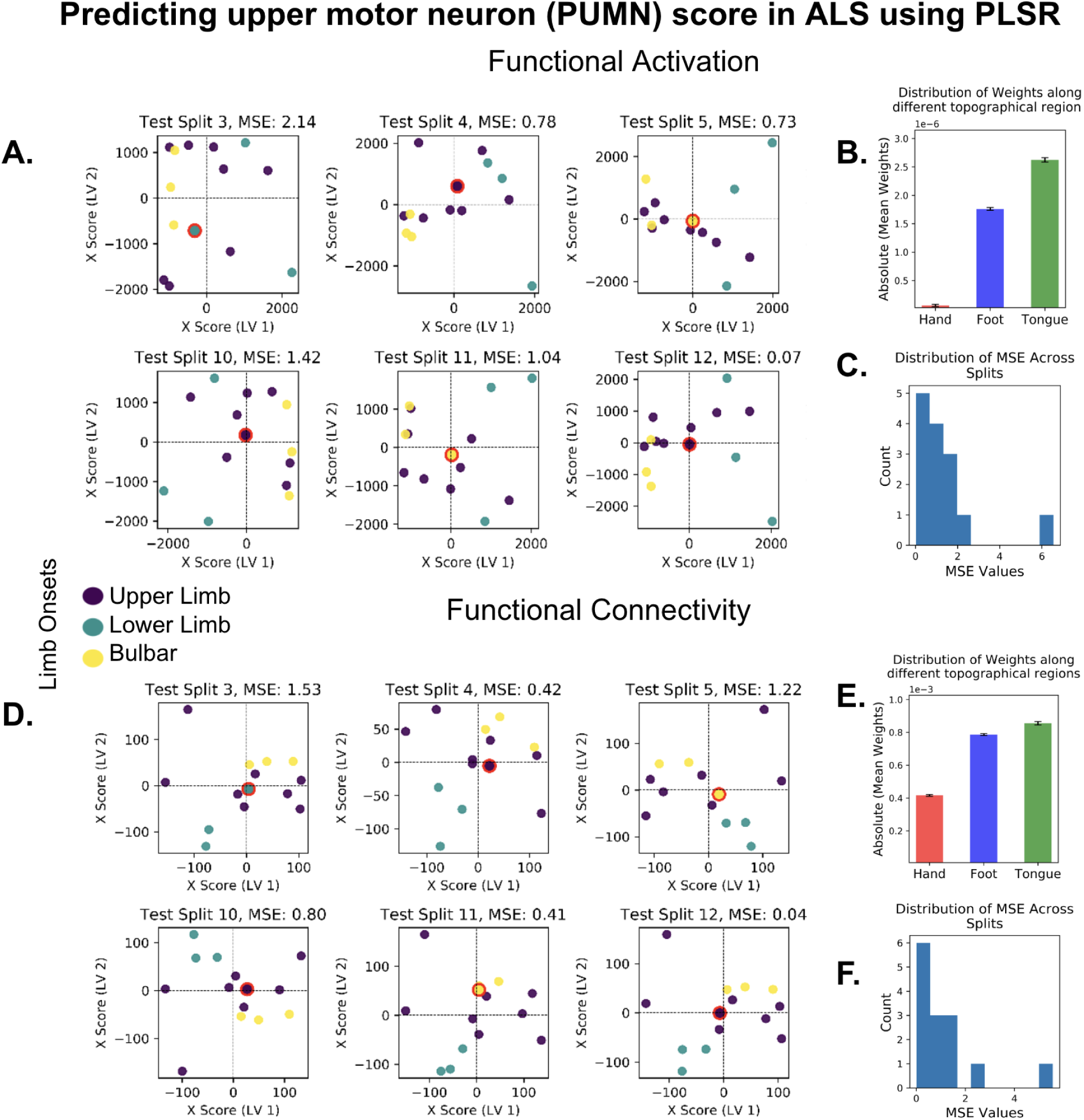
Prediction of ALS onset site based on fMRI data from sensorimotor cortex during body part movements. **(A)** + **(D)** Scatter plots depict the relationship between the first latent variable (LV1) and the second latent variable (LV2) for test subjects using PLSR, with different colored circles representing the ALS onset classes (purple: upper limb onset, cyan: lower limb onset, yellow: bulbar onset). The position of each dot in the latent space represents an individual subject, with the color indicating the onset class and the size reflecting the group distribution. Red-bordered dots highlight the test subjects. The Mean Squared Error (MSE) values for each test split are shown in the subplot titles, indicating the accuracy of prediction. **(A)** Projections for functional activation across 6 test splits, with MSE values provided for each split. **(B)** Bar plot summarizing the distribution of weights across topographical regions (hand, foot, tongue) for functional activation. **(C)** Histogram displaying the distribution of MSE values for functional activation splits. **(D)** Projections for functional connectivity across 6 test splits, with MSE values provided for each split. **(E)** Bar plot summarizing the distribution of weights across topographical regions (hand, foot, tongue) for functional connectivity. **(F)** Histogram displaying the distribution of MSE values for functional connectivity splits.

For functional connectivity, the overall MSE was 1.22 ± 0.43. The scatter plots in **Figure 3D** show better clustering of onset types compared to the functional activation, particularly for lower limb versus bulbar onset along LV1. Unlike functional activation, upper limb onset is also more clearly separated in many of the splits (e.g., splits 4, 5, 10, and 12), resulting in a more distinct grouping of onset types in the latent variable space. This is reflected in the weights calculated per topographic area, where the hand region here shows higher weights ((0.45 ± 0.1e-3). Highest weights, however, were still obtained from the tongue/face region (0.86 ± 0.1 e-3), followed by the foot region (0.79 ± 0.1 e-3, see **Figure 3E**). The histogram in **Figure 3F** shows a consistent spread of MSE values, with most values concentrated between 1 and 3, and fewer outliers compared to functional activation. Together, these results show better and more consistent performance for functional connectivity-based classification compared to functional activation-based classification of onset site across splits.

#### 3.2.2 Disease Severity Prediction

With respect to functional activation, the overall MSE is 1.22 ± 0.33 across all cross-validation splits. However, the model shows fluctuations in the LV space with respect to the projection of the subjects (see **Figure 4D**). This indicates that there is instability in the model, even though there is consistent grouping observed across all the splits. Consequently, this model was not investigated further.

**Figure 4:**
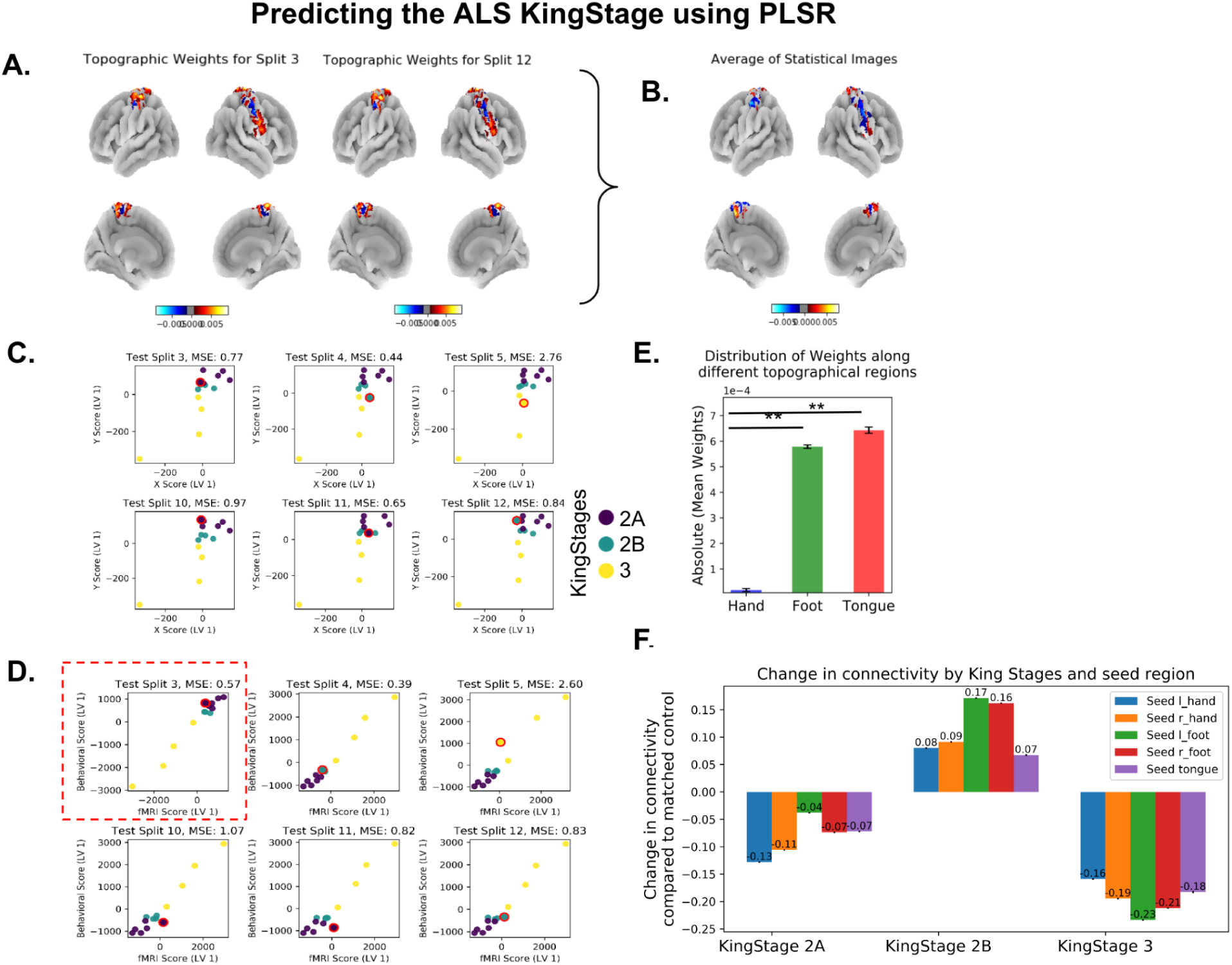
Predicting ALS King stage based on fMRI data of sensorimotor cortex during body part movements: **(A)** Topographic weights for selected splits: Inflated cortices illustrate topographic distribution of weights derived from PLSR (for example test splits 3 and 12). Color bars indicate weight values, with blue representing negative weights and red representing positive weights. These weights highlight cortical areas most associated with the ALS King stage for each test split. **(B)** Aggregated brain images show the average topographic weights across all test splits. This summary representation highlights the consistent brain regions associated with ALS progression, as indicated by the color bar where blue gradient indicate negative weights (i.e. if the connectivity in these regions increase the ALSFRS scores decrease) and more red/yellow values indicate positive weights (i.e. increased connectivity patterns in this region relates to better ALSFRS scores). **(C)** Scatter plots show the relationship between the predicted Y scores (disease stages) and the observed X scores (C - ECM, D - Brain activation) in the latent variable (LV) space for various test splits. The points with red circles represent the test subjects projected in the LV space. The color coding of the points corresponds to the King stages: 2A (purple), 2B (blue), and 3 (yellow). The mean squared error (MSE) values for each test split are indicated, reflecting the reliability of the prediction model. **(E)** Bar graph depicts the mean weights distributed across different topographical regions (hand, foot, tongue/face). Error bars represent the standard error of the mean weights. The foot and tongue/face regions show higher mean weights compared to the hand region, indicating a stronger association with ALS King’s College stages. **(F)** Bar graph shows the averaged change in functional connectivity compared to matched controls across different ALS disease stages (2A, 2B, 3). The connectivity changes are shown for seed regions corresponding to left hand, right hand, left foot, right foot, and tongue movements.

With respect to functional connectivity, the averaged MSE is 1.40 ± 0.30 across all cross-validation splits. The model shows consistent performance in predicting ALSFRS scores from functional connectivity, i.e., EC (MSE range between 0.33 and 4.60 with four splits at MSE > 1, which indicates lower predictive power). Specifically, ECM LV1 reveals distinct groupings of participants according to their ALS King stages (2A, 2B, and 3). Participants in ALS King stage 3 are consistently grouped below ‘0’ on the y-axis, while participants in ALS King stages 2A and 2B are grouped above ‘0’ on the y-axis. This consistent grouping across different cross-validation splits suggests that the first latent variable of the functional connectivity profile effectively captures variability associated with ALS disease severity (see **Figure 4**). When projecting the test subject’ data onto the latent variable space, data points fall within the same grouped regions as the training data points observed in the scatter plot (see **Figure 4 C**, red outlined points). This confirms the model’s ability to generalize and predict ALS disease stages based on connectivity (ECM) values, which has been less successful based on functional activation (see above).

To explore this effect further, **Figure 4F** illustrates the averaged changes in functional connectivity across different ALS disease stages compared to matched controls, which presents as an inverted u-shaped profile: In King stage 2A, functional connectivity is decreased across all seed regions relative to controls, with reductions observed in all regions (left hand: -0.13, right hand: -0.11, left foot: -0.04, right foot: -0.07, and tongue: -0.07). In King stage 2B, an increase in connectivity can be observed in patients versus controls again for all regions (left hand: +0.08, right hand: +0.09, left foot: +0.17, right foot: +0.16, and tongue: +0.07). In King stage 3, connectivity is again reduced in patients versus controls in all regions left hand: -0.16, right hand: -0.19, left foot: -0.23, right foot: -0.21, and tongue: -0.18). See **Supplementary** Figure 1 and **Supplementary Table 1** for longitudinal data sampled from different time points available for n=3 patients.

In summary, with respect to (1) and (2), we have shown that BOLD signal change in sensorimotor cortex during body part movements can be used to successfully classify a group of participants into patients versus controls, to classify patients with respect to disease severity and site of onset, and that functional connectivity is a better predictor for both disease onset and disease severity compared to functional activation. We further show that functional connectivity changes in patients versus controls present with an inverted u-shaped profile, with reduced connectivity in early disease stages, higher connectivity in middle disease stages, and a drop in connectivity in more severe disease stages.

### 3.3. Topographic versus atopographic profile

Finally, we investigated if (3) disease-defining functional information is specific to the first-affected topographic location in the sensorimotor cortex (i.e., topographic, reflecting the iron accumulation) or not specific to the first-affected topographic location in the sensorimotor cortex (i.e., atopographic, reflecting the calcium accumulation), and whether there is any topographic location in MI that is highly predictive of the severity of the disorder irrespective of onset site. Above, we have reported better classification accuracies in the patient versus control classification in regions of the behaviorally non-first affected body part compared to regions of the first-affected body parts (see section 3.1.). This already shows that functionally disease-defining regions in MI are not strongest in the area that is behaviorally first-affected.

In addition, we have reported higher weights in the foot and tongue regions compared to the hand region for the prediction of disease onset area. To explore this effect of potentially more predictive areas in MI further, we computed the model’s weight distribution (based on connectivity) of the King’s Stage classification and projected them back into voxel space. In this way, the regions that drive the classification for disease severity can be identified precisely in MI. Results reveal that the highest mean weights are present in the foot and tongue/face regions, with the hand region showing marginally low weights (hand = 1.72e-05 ± 5.78e-06, foot = 5.7e-04 ± 6.828e-06, face = 6.4e-04 ± 1.22e-05, see **Figure 4E**). A paired t-test between these weight distributions indicates a significant difference in weight distributions between hand vs. foot region (t(13) = -8.90, p = 6.84e-07), and hand vs. face region (t(13) = -3.69, p = 2.72e-03), and no significant difference between foot and face region (t(13) = -0.49, p = 6.34e-01) (see **Figure 4E**). This indicates that higher connectedness (i.e., centrality) in the face and lower limb regions within the brain’s network is predictive of the King stage of the disease, whereas this is less the case for the hand region. Both for the prediction of disease onset and disease severity, the foot and face areas show hence the highest predictive power, whereas the hand area shows lowest predictive power.

## 4. Discussion

ALS is a rapidly progressing neurodegenerative disease characterized by the loss of motor control and associated changes in the cytoarchitectural architecture of MI (Al-Chalabi et al., 2016). Whereas previous studies mostly focused on structural alterations, the present study aimed to uncover the distinct functional activity signature of MI associated with ALS onset site and progression using ultra-high field 7T-fMRI, rSRM, and PLS regression analysis. Our results (1) confirm that participants can be successfully classified into ALS-patients and controls based on their BOLD signal change in sensorimotor cortex recorded during body part movements, (2) show that latent variables specifically of the functional connectivity maps (but less so of the functional activation maps) reflect disease severity and disease site of onset, and (3) demonstrate that disease-defining functional information is not specific to the first-affected topographic field in sensorimotor cortex, but that the foot and face areas in MI are most predictive of the severity of the disorder irrespective of onset site. The results of our study contribute to a deeper understanding of the cortical mechanisms underlying ALS, and offer a novel methodological approach to characterize the disease in future clinical studies.

We first show that ALS patients can be successfully distinguished from healthy controls based on both functional activation and connectivity patterns in the sensorimotor cortex, with an overall classification accuracy of 0.91 ± 0.26. This aligns with previous studies showing distinct neural activity patterns in ALS patients, particularly in the motor regions, which are known to be affected early in disease course (e.g., Rubio et al., 2022; Trojsi et al., 2012; Xie et al., 2023). The clear separation observed in the latent variable space supports the overall idea that ALS progression can be effectively tracked through neuroimaging markers, potentially aiding in early diagnosis and personalized treatment strategies.

Both the prediction of disease onset site and severity is more effective using functional connectivity rather than functional activation maps. This is an important insight as it indicates that specifically the networks co-activated with voluntary movement control in ALS are most predictive of disease stage, rather than the net activation in MI during movements as such. Prior studies have indicated that the functional connectivity profile in ALS patients is predictive of disease stage, however, those studies often focused on cognitive decline and associated network changes (De Marchi et al., 2021), or investigated resting state networks only (Metzger et al., 2024). The further result that functional connectivity changes are decreased compared to controls in King stage 2A, are increased in King stage 2b, and are decreased again King stage 3 provides a further indication that functional network changes in MI are highly sensitive to disease progression (Notturno et al., 2023). This aligns with a study using MEG recordings from early-diagnosed ALS patients during speech tasks, which found greater sensory correlations and beta band connectivity in ALS patients compared to healthy controls (Dash et al., 2024). This suggests that ALS patients exhibit increased functional connectivity in early stages of the disease to compensate for an initial decline in connectivity by recruiting additional networks as support architecture. The later decline in connectivity could then signal more severe disease stages with more pronounced myelin loss, where this compensation strategy fails. For translational applications, this could imply that focusing on maintaining this initial compensation strategy at a high level, i.e., focusing on preventing the decline in connectivity that is observed at later disease stages, may be a strategy to slow down disease progression. This finding is also in line with existing literature (e.g. van den Bos et al., 2019) that suggests early compensatory mechanisms in ALS, where the brain attempts to maintain motor function despite ongoing neurodegeneration.

With respect to the a/topographic functional architecture of ALS, our definition of ‘topography’ is based on earlier structural investigations of the same patients, which revealed that the first-affected body part shows higher iron accumulation in deep layers of MI compared to the later-affected regions in MI (Northall et al. 2024). With this definition, a ‘topographic’ functional signature would show disease-defining features specifically in earlier-affected compared to later-affected regions of MI, following the deep layer iron accumulation. On the other hand, an atopographic functional signature could also be driven by stronger functional changes in newly-affected areas of MI, and would indicate that disease-defining functional features are not necessarily most pronounced in the most clinically-affected body part. This assumption is based on the observation that the accumulation of nQSM in ALS (potentially reflecting calcium) is specifically pronounced in low-myelin borders of MI, which are located between body part areas of MI (Northall et al. 2024), and that we detected generally lower myelination of the face area with aging, indicating vulnerability of this region (Northall et al. 2023). Our investigation reveals that regions first affected by ALS exhibit lower group (i.e., ALS vs. control) classification accuracy compared to non-first affected regions in MI when using functional activation time series. This suggests that functional activation in the first-affected regions may not fully capture the early pathological changes driving ALS progression. Interestingly, the stronger classification accuracy observed in non-first affected regions likely reflects the progression to second or third affected body parts, as indicated by higher King stages. These non-first affected regions may exhibit more pronounced functional alterations, potentially due to advanced pathology resulting from later disease stages. The higher classification accuracy in these regions shows the importance of considering functional activation patterns over time as ALS progresses.

In contrast, the structural changes observed in previous studies, such as higher iron accumulation in the deep layers of MI in first-affected regions in these patients (Northall et al. 2024), suggest that early ALS pathology is better captured through structural imaging rather than functional activation. Instead, the functional activity profiles we observed may align more closely with atopographic calcium accumulation in MI, which has been shown to predict later demyelination. However, our findings are limited by the absence of direct data on spinal cord and brainstem involvement or second motor neuron degeneration, which could significantly impact both functional activation and connectivity patterns. Future studies incorporating these regions and combining functional and structural markers may provide a more comprehensive understanding of ALS progression. Critically, the calcium accumulation in patients was specifically high in the low-myelin borders in MI that separate distinct body part areas (Kuehn et al. 2017) and that are assumed to be connected to different, non-topographic functional networks compared to the body part areas in MI (Gordon et al. 2023). This pattern of results hence indicates that disease-defining network changes do not necessarily overlap with the earliest-affected body parts and the hotspots of iron accumulation. In order to successfully track disease progress and to investigate the effect of medication, an accompanying mapping of functional network changes is therefore to be recommended based on our results. In addition, if this is confirmed by future studies involving more patients, a potential novel mechanistic target would be activating/maintaining the low-myelin borders in MI to slow down disease progression.

We also show that both disease onset site and severity can best be predicted by the foot and face areas in MI, rather than the hand area. The significant differences in weight distribution across different regions of the primary motor cortex further support this finding. This regional specificity could be due to the differential vulnerability of motor neurons in these regions to ALS-related pathologies such as iron accumulation (Donatelli et al., 2022; Kwan et al., 2012). In a recent study, (Northall et al., 2023b) found that the hand region in MI in older adults shows less age-related iron accumulation compared to regions representing other body parts, arguing that the hand region is potentially less vulnerable compared to other areas in MI to effects of aging (Northall et al., 2023b). This resilience could explain why the hand region displays fewer functional alterations, thus making it a less effective marker for classification in ALS. Another possibility is that hand dysfunction is detected earlier behaviorally due to the very high level of hand dexterity in humans. This finding is also consistent with the fact that bulbar impairment is linked to shorter survival in ALS (Stegmann et al., 2020). More pronounced changes in bulbar activity could serve as a more sensitive marker for overall disease aggressiveness and progression compared to other body parts.

## 5. Conclusion

In conclusion, this study demonstrates the potential of using rSRM and PLSR in combination with 7T-fMRI for uncovering the functional correlates of ALS progression and onset even in small patient samples. Our findings provide critical insights into the disease’s atopographic patterns, and highlight the importance of the face/tongue and foot regions in disease assessment, hinting towards their potential as biomarkers for tracking disease onset and progression. These results pave the way for future research aimed at improving the diagnosis and management of ALS through advanced neuroimaging techniques.

## 6. Acknowledgement

This work was funded by the Deutsche Forschungsgemeinschaft (DFG) (MA 9235/3-1/SCHR 1418/5-1 (501214112)), SFB 1436 Z02 (Project-ID 425899996), and Else Kröner Fresenius Stiftung: 2019-A03

## 7. Data availability

fMRI/MRI data will be made available upon request

## 8. Code availability

https://github.com/avinashkalyani/ALS_PLSr

## 10. Credit authorship contribution statement

**Avinash Kalyani:** Conceptualization, Investigation, Formal analysis, Methodology, Writing - original draft. **Alicia Northall:** Data collection, Formal analysis, Methodology, Writing - review, and editing. **Stefanie Schreiber:** Conceptualization, Writing - review, and editing. **Jascha Brüggemann:** Data collection. **Stefan Vielhaber:** Data collection. **Marwa Al Dubai:** Data collection. **Abrar Benramadan:** Data collection. **Oliver Speck:** Writing - review, and editing. **Hendrik Mattern:** Writing - review, and editing. **Christoph Reichert:** Supervision, Conceptualization, Methodology, Writing - review, and editing. **Esther Kuehn:** Supervision, Investigation, Conceptualization, Methodology, Writing - review, and editing.

