## Supplemental Material for "Individualized Phenotyping of Functional ALS Pathology in Sensorimotor Cortex"

### Supplementary Material

#### S.1 Longitudinal percent signal changes exceed the change in connectivity in ALS

To investigate the progression of brain activity over time in ALS patients, we measured both percent signal change and change in Eigenvector Centrality Mapping (ECM) values at different time points for three subjects: P4 (with data at 11 months and 3 years), P1 (with data at 5 months), and P6 (with data at 7 months). As illustrated in Figures 5A and 5B, these measures varied across different time points for each patient, reflecting the dynamic nature of ALS. In P4, the percent signal change in the hand and tongue regions initially increased over 11 months but then significantly decreased within 3 years, while the foot region showed a continuous decline over the same period. Despite these changes in functional activation, ECM values remained relatively stable across all regions, indicating that while neural activity fluctuated, the overall network connectivity did not exhibit substantial alterations. For P1, there was a steady increase in percent signal change across all regions (hand, foot, and tongue) after the 5-month period. ECM values for the hand and foot regions remained stable, but there was a slight decrease in the tongue region, suggesting localized changes in connectivity that did not correspond to broader network disruptions. P6 demonstrated more variability, with percent signal changes showing a decrease in the hand region, stability in the foot region, and an increase in the tongue region over 7 months. ECM values for P6 remained consistent across all regions, indicating that the fluctuations in functional activation did not translate into significant changes in overall connectivity (see **Table 2** for values).

These observations highlight the variability in functional activity and connectivity in ALS patients over time. The analysis suggests that changes in functional activation, as indicated by percent signal change, are more pronounced than changes in connectivity, as measured by ECM values. This aligns with current understanding in ALS research, where compensatory increases in neural activity are often observed in the early stages of the disease, followed by declines as the disease progresses and motor neuron degeneration becomes more extensive. This pattern underscores the importance of monitoring both functional activation and connectivity to fully understand the progression of ALS.

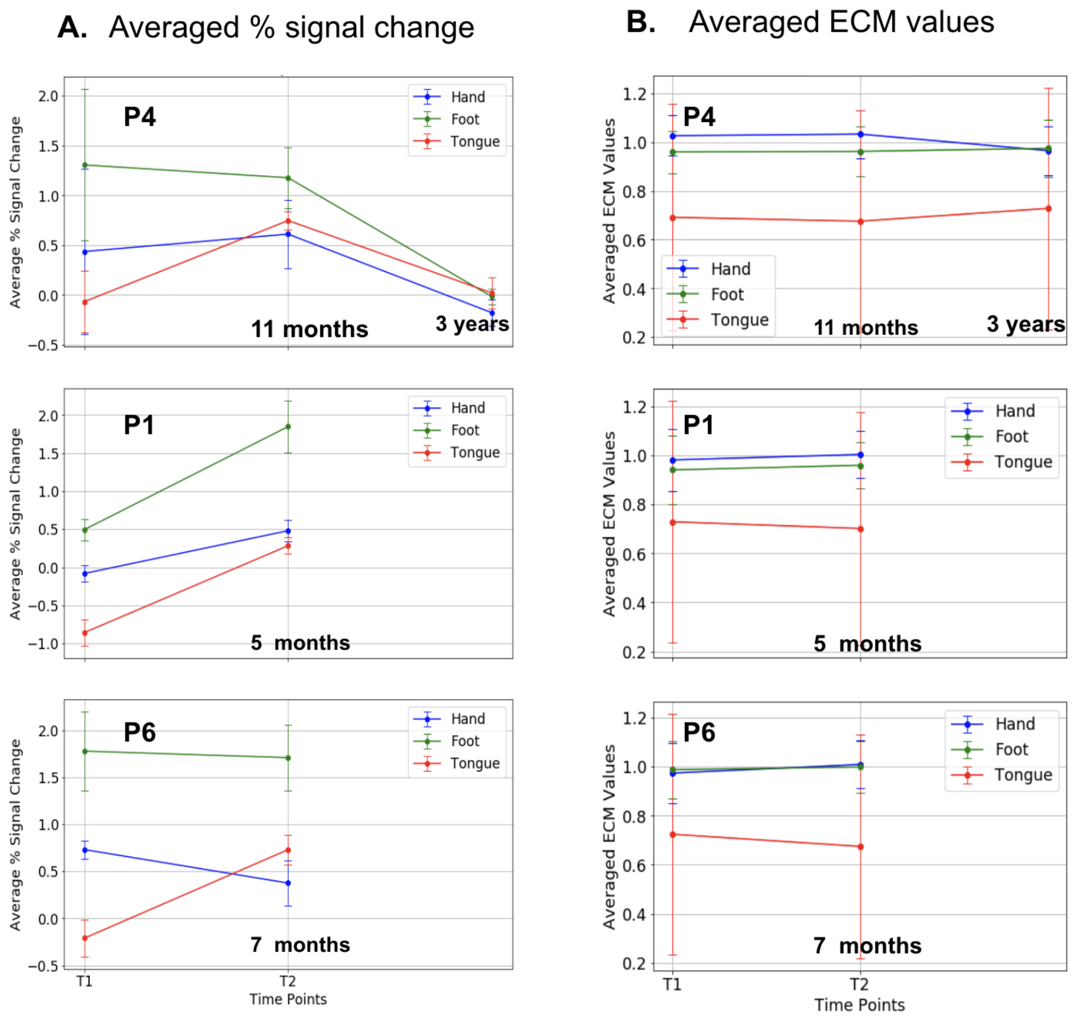


#### Clinical assessment scores overtime

| Patients | T1 | T2 | T3 |
|---|---|---|---|
| **P4** | ALSFRS-R Total: **37** (bulbar: 12, fine : 6, gross : 7, breathing: 12), KS: **2B** | ALSFRS-R Total: **36** (bulbar: 12, fine : 6, gross : 7, breathing: 11), KS: **2B** | ALSFRS-R Total: **41** (bulbar: 11, fine : 10, gross : 9, breathing: 11), KS: **2B** |
| **P1** | - | ALSFRS-R Total: **44** (bulbar: 12, fine : 9, gross : 11, breathing: 12) KS: **2A** | - |
| **P6** | ALSFRS-R Total: **41** (bulbar: 9, fine : 9, gross : 11, breathing: 12) KS: **2B** | ALSFRS-R Total: **38** (bulbar: 9, fine : 7, gross : 10, breathing: 12) KS: **3** | |

**Supplementary Figure 1. Changes in % signal change and functional connectivity over time: (A)** Averaged % signal change and **(B)** averaged Eigenvector Centrality Mapping (ECM) Values: Line graphs illustrate averaged % signal change and ECM values in brain activity for three subjects across different time points: +11 months and +3 years after the first san for P4, +5 months after the first scan for P1, +7 months after the first scan for P6. Data is extracted from hand (blue), foot (red), and tongue (green) regions in MI, highlighting the variations in functional activation and connectivity over time. The table at the bottom row shows the behavioral ALSFRS-R scores measures over time, and the King Stage (KS) for the patients. Note: P1 was not recorded at the first time point

**Supplementary Table 1 Percent signal change and connectivity changes in ALS patients over time**

| Patient | Region | Time Point | Percent Signal Change (mean ± SEM) | ECM Values (mean ± SEM) |
|---|---|---|---|---|
| P4 | Hand | Baseline | 0.41 ± 0.83 | 1.03 ± 0.08 |
| | | 11 months | 0.58 ± 0.34 | 1.03 ± 0.10 |
| | | 3 years | 0.18 ± 0.13 | 0.97 ± 0.12 |
| | Foot | Baseline | 1.26 ± 0.76 | 0.96 ± 0.09 |
| | | 11 months | 1.21 ± 0.31 | 0.96 ± 0.10 |
| | | 3 years | 0.04 ± 0.08 | 0.96 ± 0.10 |
| | Bulbar | Baseline | 0.02 ± 0.31 | 0.69 ± 0.46 |
| | | 11 months | 0.74 ± 0.09 | 0.68 ± 0.46 |
| | | 3 years | 0.03 ± 0.16 | 0.73 ± 0.50 |
| P1 | Hand | Baseline | 0.15 ± 0.11 | 0.98 ± 0.13 |
| | | 5 months | 0.38 ± 0.14 | 1.00 ± 0.09 |
| | Foot | Baseline | 0.37 ± 0.14 | 0.94 ± 0.14 |
| | | 5 months | 1.74 ± 0.34 | 0.96 ± 0.09 |
| | Bulbar | Baseline | -0.89 ± 0.17 | 0.73 ± 0.49 |
| | | 5 months | 0.25 ± 0.11 | 0.70 ± 0.48 |
| P6 | Hand | Baseline | 0.86 ± 0.10 | 0.97 ± 0.12 |
| | | 7 months | 0.39 ± 0.24 | 1.01 ± 0.10 |
| | Foot | Baseline | 1.80 ± 0.42 | 0.99 ± 0.12 |
| | | 7 months | 1.82 ± 0.35 | 1.00 ± 0.11 |
| | Bulbar | Baseline | 0.16 ± 0.20 | 0.72 ± 0.49 |
| | | 7 months | 0.75 ± 0.16 | 0.67 ± 0.46 |
